# A cross-attention transformer encoder for paired sequence data

**DOI:** 10.1101/2023.12.11.571066

**Authors:** Ceder Dens, Kris Laukens, Pieter Meysman, Wout Bittremieux

## Abstract

Transformer-based sequence encoding architectures are often limited to a single-sequence input while some tasks require a multi-sequence input. For example, the peptide–MHCII binding prediction task where the input consists of two protein sequences. Current workarounds to solve this input-type mismatch lack resemblance with the biological mechanisms behind the task. As a solution, we propose a novel cross-attention transformer encoder that creates a cross-attended embedding of both input sequences. We compare its classification performance on the peptide–MHCII binding prediction task to a baseline logistic regression model and a default transformer encoder. Finally, we make visualizations of the attention layers to show how the different models learn different patterns.

## Introduction

Recently, transformers have proven to be very valuable for natural language processing. BERT [1] (Bidirectional Encoder Representations from Transformers) is an architecture based on the encoder of a transformer and was successfully applied to various tasks like text summarization, intention detection and topic classification. Recently, its potential for analyzing protein sequence data was shown [2]–[4]. One of the limitations of these BERT models for protein sequences is that they are designed to only process a single protein sequence at once, making them less suitable for e.g. contact and interaction prediction tasks, because they take two protein sequences as input. Multiple workarounds to allow these kinds of predictions exist but lack resemblance with the biological mechanisms behind the task. A possible approach is to concatenate both input sequences and use this as a single input for the model [5]. A possible problem with this approach is that it might be hard for the model to learn the intrinsic features and properties of both sequences and also learn the cross-sequence patterns that determine the final output prediction (e.g. interaction). A second possible workaround is to have a separate BERT model for both sequences [6], concatenate the outputs of these models at the end, and send them through the last part of the model (e.g. a multi-layer perceptron) to make the final prediction. The downside of this approach is that there are no transformer layers used on both sequences together, while the attention mechanism is great at learning these inter-token relations and patterns. The last workaround that is often used for dual-input transformer models is a cross-attention mechanism. The traditional BERT architecture only uses self-attention layers, in each of these layers the attention is calculated using the input sequence twice (hence self-attention) and afterwards this attention is again applied to the original input embedding to get the self-attended output embedding. Using cross-attention instead, the attention is calculated between both input sequences and is then applied to only one of the input embeddings with as result a cross-attended embedding for the other sequence as output. As a consequence, the output of such a cross-attention layer is an embedding for only one of the two input sequences. This approach can be very useful if you have one ‘main’ input and want to add information from a second input. But, in the case of e.g. interaction prediction, we would prefer to have a cross-attended embedding for both sequences that treats them both equally important.

As an alternative, we present a new cross-attention layer that does produce a cross-attended embedding of both inputs as output. This layer can be used in combination with concatenated self-attention layers and parallel self-attention layers. We test multiple model architectures with and without cross-attention layers on the peptide–Major Histocompatibility Complex Class II (peptide–MHCII) binding prediction task. An MHCII molecule is a protein complex on the cell surface of antigen presenting cells that presents peptides. The task at hand is to predict which MHCII molecules are able to present which peptides. A peptide is a short protein sequence and the MHCII protein complex can also be represented as a relatively short protein sequence. This makes this a suitable input for our method.

## Methods

### Cross-attention layer

Our new cross-attention (CA) layer is similar to the regular self-attention layer. But, we made two major changes: the inputs used to calculate the Query, Key and Value matrices are different and we project the output of the attention calculation to new dimensions (fig. 1). The attention matrix is always calculated between the Query and Key. In the case of self-attention, both Q and K would be derived from the same input. For our cross-attention layer, we use one of the sequences to create the Query and the other sequence to create the Key, this results in an attention matrix with dimensions len(s_a) x len(s_b). Normally, cross-attention would then apply this attention matrix to one of the two inputs, resulting in an embedding for the other input. Our cross-attention layer aims to create a cross-attended embedding for both inputs, that is why we use the concatenated inputs to create the Value matrix. As a result, the attention matrix will be applied to both inputs. For this to work, we transform the cross-attention matrix to a matching shape. This transformation is implemented by applying a linear layer (from len(s_b) to len(s_a+s_b)), this output is transposed and again inputted in a linear layer (from len(s_a) to len(s_a+s_b)). The projected cross-attention matrix has size len(s_a+s_b) x len(s_a+s_b), multiplying this with our Value vector results in a cross-attended embedding for both sequences.

**Figure 1.**
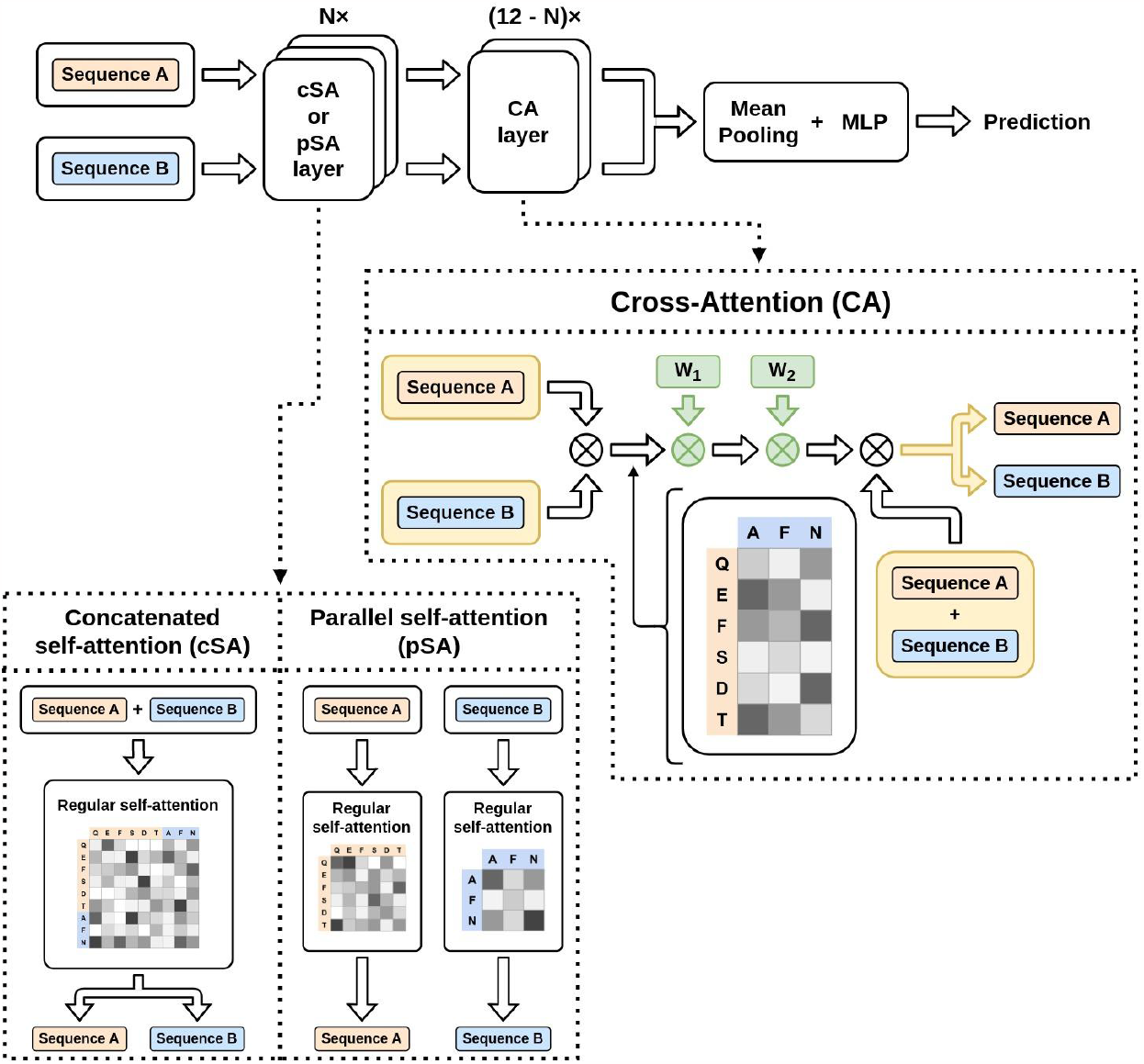
Schematic overview of the Cross-Attention BERT architecture. On top, the full model architecture, consisting of N cSA or pSA layers followed by 12-N CA layers, a mean pooling and MLP layer. Below, an overview of the difference between a cSA and pSA layer and a detailed schematic of the CA layer. The CA layer starts with multiplying sequence A and B, this results in the cross-attention matrix. Then this is again multiplied with new weights W1 and W2 to transform the cross-attention matrix to its new dimensions. Finally this is multiplied again with concatenated sequences A and B to result in the cross-embedded output for sequence A and B.

### Parallel self-attention layer

As both inputs can have different properties and mechanisms, it is not always desirable to apply the self-attention mechanism on the concatenated inputs. On top of that, the interactions between the inputs should be learned in the cross-attention layer(s). That is why we use parallel self-attention (pSA) layers. This layer applies regular self-attention on both inputs and keeps them completely separate (see Fig. 1). This makes that these layers focus on the structure and patterns of both inputs separately.

### Final model architecture

As shown in figure 1, the final model starts with N cSA or pSA layers. All our comparisons were done with N being 10, 11, or 12. Then 12 - N CA layers follow, meaning that there can be 2, 1 or 0. Finally, a mean pooling and a multilayer perceptron (MLP) is applied to get the output prediction.

### Finetuning pretrained model

All our models are only finetuned, we use the pretrained weights from the TAPE model [2], which is pretrained on a large collection of protein sequences, to initialize our layers. The layers are always initialized with the weights of the corresponding pretrained layers, meaning that our Nth layer is initialized with weights from the Nth pretrained layer. The cSA layers can be directly initialized with the corresponding pretrained layers. For the pSA layers, both parallel layers are initialized with the weights of the same pretrained layer. Most of the weights of the CA layers are also initialized with the pretrained weights, only the two linear layers we added are not initialized.

### Cross-validated hyperparameter selection

To find the best model, we performed 5-fold cross-validation. Each configuration was first trained and afterwards tested on the validation dataset, the configuration with the best validation ROC-AUC was chosen. We tested different values for the learning rate (0.1, 0.01, 0.001, and 0.0001) and optimizer (SGD and AdamW) and experimented with different layer combinations. All models have 12 layers in total, we experimented with using 12 pSA layers, 12 cSA layers, 10 or 11 pSA layers followed by 1 or 2 CA layers and 10 or 11 cSA layers followed by 1 or 2 CA layers. At the end, 3 models were chosen based on the best parameter configuration. The best model using cSA and CA layers, the best model using only cSA layers and the best model using pSA and CA layers.

### Logistic regression model

As a baseline model, we implemented a logistic regression on an embedding of the pretrained TAPE model [2]. We have two versions of this model, a concatenated and parallel version. The concatenated version calculates an embedding from the concatenated sequences using the frozen pretrained model, this embedding then goes through a single linear layer and sigmoid activation function. The parallel version calculates an embedding for both input sequences from the frozen pretrained model separately, then these embeddings are concatenated and again go through a single linear layer and sigmoid activation. All embeddings are calculated with the frozen default TAPE BERT model.

### Peptide–MHC data

Our model is evaluated on peptide–MHCII binding data. The peptide and MHC are both relatively short protein sequences. The task is to predict whether a given MHC would bind/present a given peptide. The same dataset as BERTMHC [5] was used to train and test our model. The train and validation data was collected from IEDB [7] up to 2016 and 5 train/val cross-validation splits were made with minimal overlap between the train and validation data. The test data is an external dataset consisting of IEDB data after 2016 and an independent dataset from the Dana–Farber repository [8]. All samples already present in the train/val data were removed.

### Visualization

Visualizations of the (self-)attention matrices were made with the python package bertviz [9]. This package supports visualizations of cSA (regular self-attention) and CA layers. One can also make visualizations of an encoder-decoder model, this functionality was reused to create visualizations for both self-attention matrices in the pSA layers.

## Results

### Performance of logistic regression models

5-fold cross-validated hyperparameter selection was performed on the logistic regression models. The best concatenated and the best parallel logistic regression model was selected based on the validation performance. Both best models use the SGD optimizer with a learning rate of 0.1. The performance reported is the average ROC-AUC and average standard deviation over the 5-fold cross-validation. The parallel logistic-regression model has a validation ROC-AUC of 74.23% ± 2.12%, the concatenated model has a validation ROC-AUC of 73.46% ± 2.27% (fig. 2a). The test ROC-AUC of the parallel model is 58.39% ± 0.47% and that of the concatenated model is 56.39% ± 0.79% (fig. 2b). On the test data, the parallel model slightly outperforms the concatenated model.

**Figure 2:**
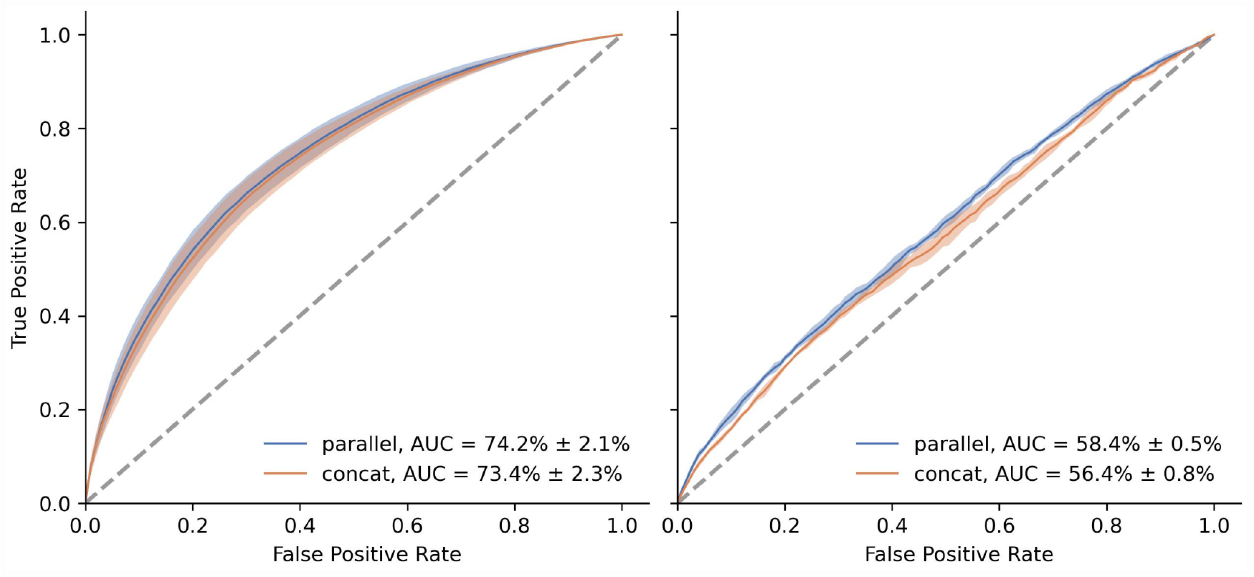
ROC-curves of (a) validation and (b) test performance of the parallel and concatenated logistic regression models. The average ROC-curve with its standard deviation is plotted.

### Performance of cross-attention models

After doing 5-fold cross-validated hyperparameter selection we selected 3 model configurations based on the validation performance: the best model containing cSA layers, the best model consisting of only cSA layers (as a reference model without CA layers) and the best model containing pSA layers. For all three models, a learning rate of 0.1 and the SGD optimizer resulted in the best validation performance. The performance reported is the average ROC-AUC and average standard deviation over the 5-fold cross-validation. The best model containing cSA layers has 10 of these followed by 2 CA layers. It has a validation ROC-AUC of 85.61% ± 1.27%. The best model with only cSA layers has a validation ROC-AUC of 85.31% ± 1.47%. The best model containing pSA layers has 11 of these followed by 1 CA layer. Its validation ROC-AUC is 84.38% ± 1.46% (fig. 3a). Testing these models on the external test set results in a ROC-AUC of 66.44% ± 0.81% for the cSA-CA model, a ROC-AUC of 66.15% ± 0.70% for the CA only model and a ROC-AUC of 64.50% ± 0.79% for the pSA-CA model (fig. 3b). The model with only cSA layers and the model with a combination of cSA and CA layers have a similar performance. On the test data, the model with a combination of pSA and CA layers performs slightly worse than the other two models.

**Figure 3.**
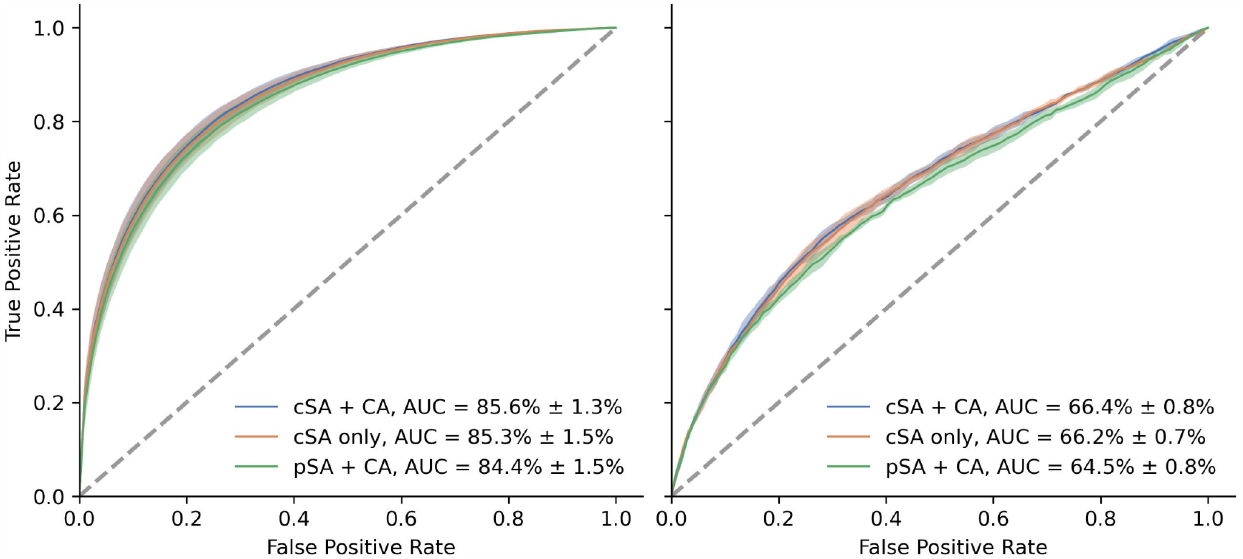
ROC-curves of (a) validation and (b) test performance of the best model for the three layer configurations. The average ROC-curve with its standard deviation is plotted.

### Visualization of different models

Visualizations of the attention matrices were made with the python package bertviz. Fig. 4 and Fig. 5 show the attention matrices of part of the layers of the cSA - CA model. Respectively 3 of the 10 cSA layers and the 2 CA layers. Fig. 6 shows the attention matrices of 3 of the cSA layers of the cSA only model. Fig. 7-9 show the attention matrices of part of the pSA - CA model layers. Respectively 3 of the 11 pSA MHC layers, 3 of the 11 pSA peptide layers, and the final CA layer.

**Figure 1.**
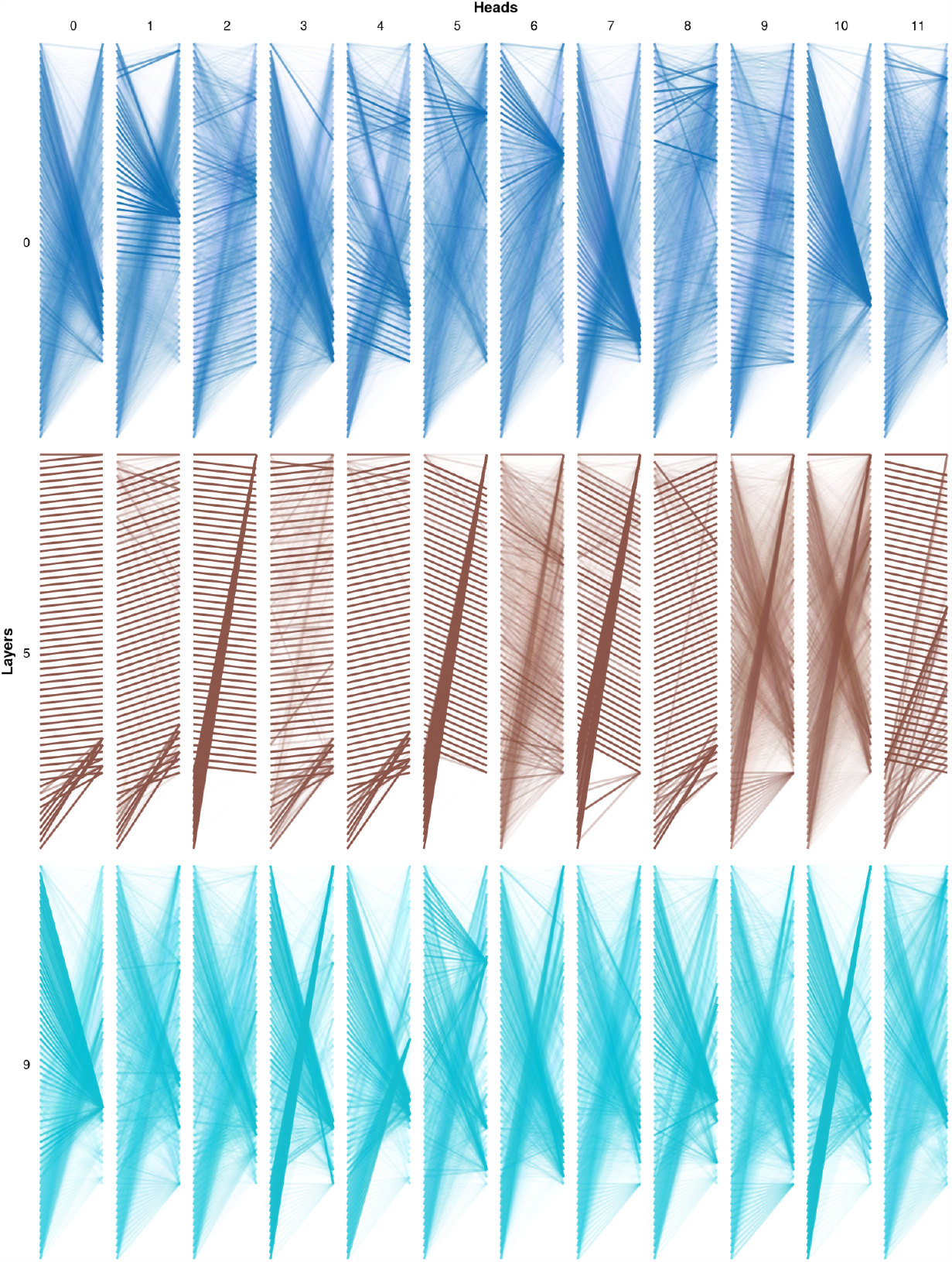
cSA-CA model cSA layer 0-9 visualization.

**Figure 2.**
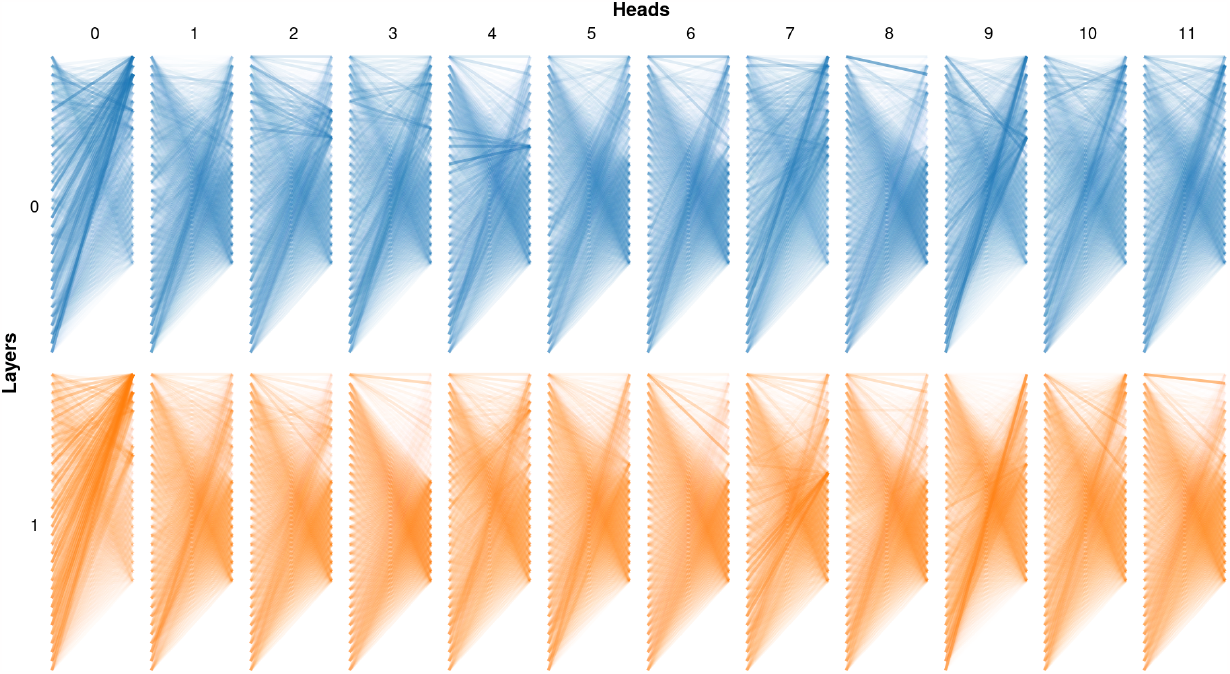
cSA-CA model CA layer 10-11 visualization.

**Figure 3.**
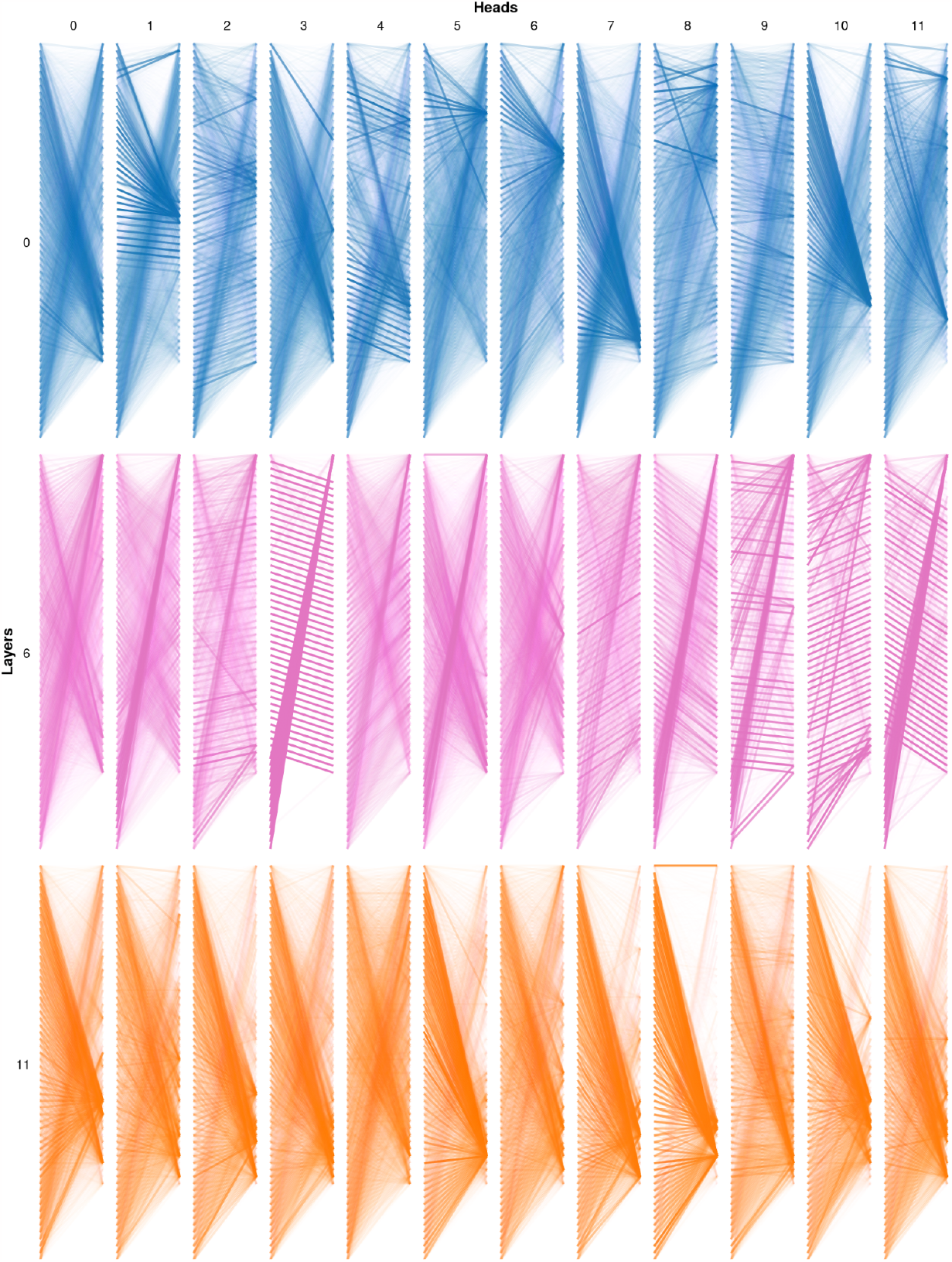
cSA only model cSA layer 0-11 visualization.

**Figure 4.**
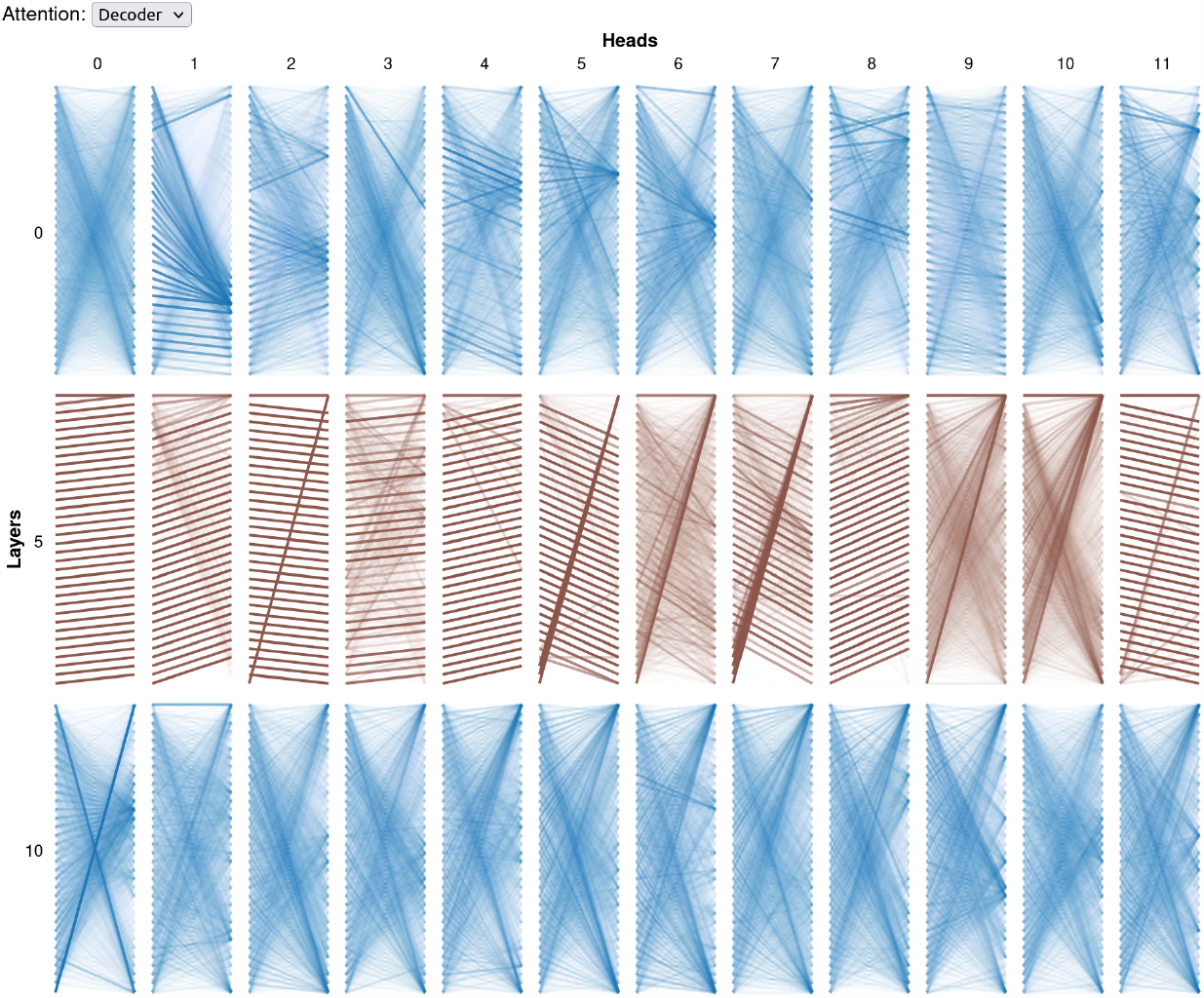
pSA-CA model pSA MHC layer 0-10 visualization

**Figure 5.**
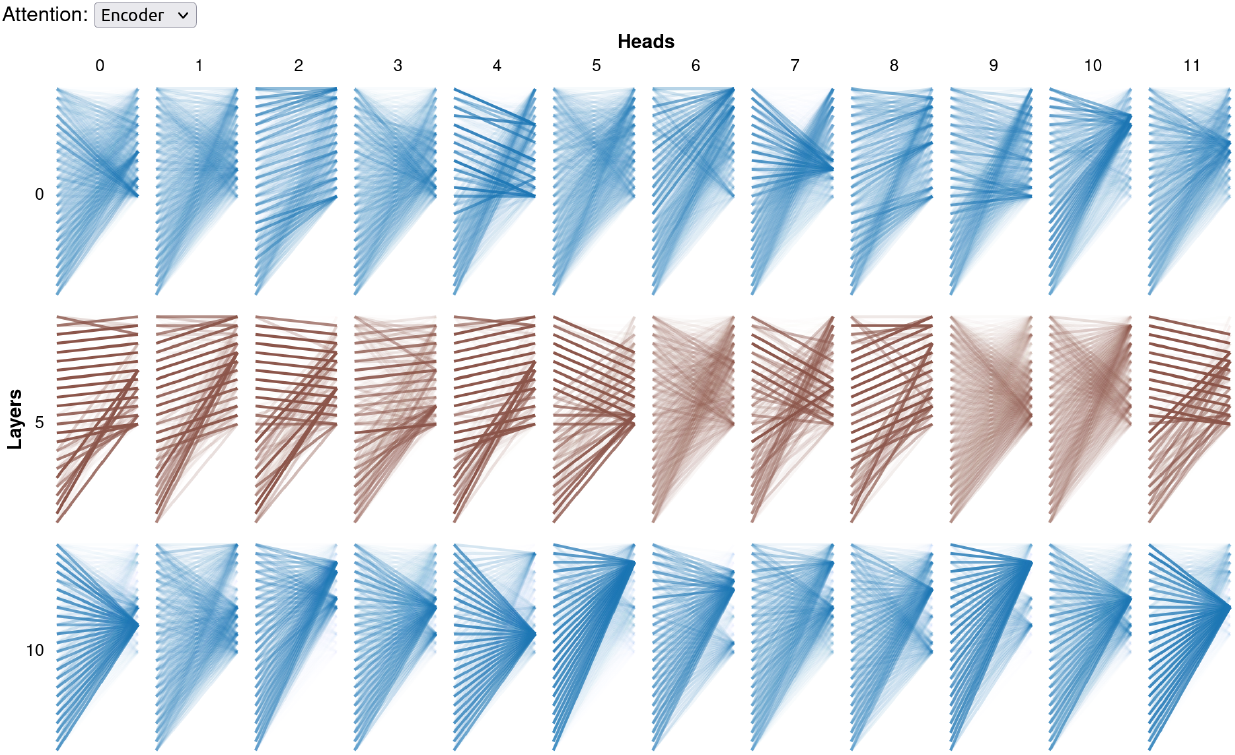
pSA-CA model pSA peptide layer 0-10 visualization

**Figure 6.**
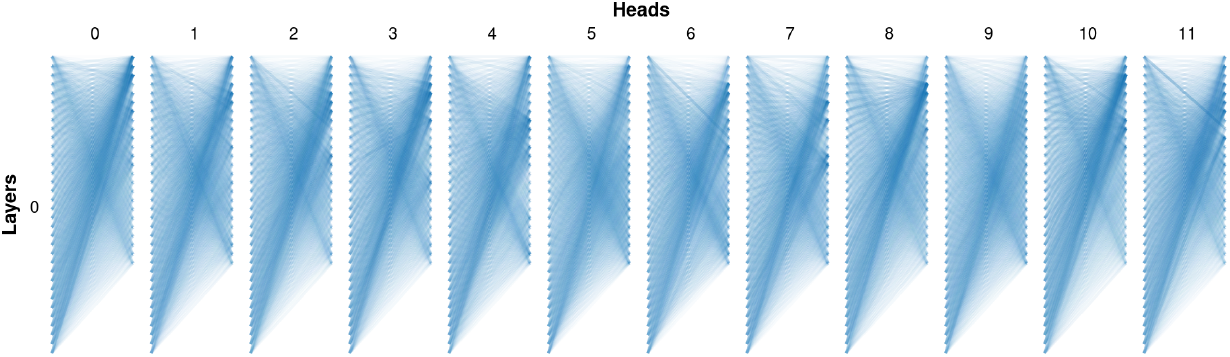
pSA-CA model CA layer visualization.

## Discussion

Current transformer models learning from a combination of two input sequences have limited resemblance with the real-world mechanisms behind the data, possibly hindering them to learn these patterns. In this work, we presented an adapted cross-attention layer that treats both input sequences equally and creates a cross-attended embedding for both sequences as output. We hypothesized that first using pSA layers that learn the intrinsic features of both sequences separately and then using CA layers to learn the combination of the two sequences would work best. Testing on peptide–MHC binding data showed that this model architecture slightly underperformed compared to using only cSA layers or a combination of cSA and CA layers. This might be due to the limited complexity of the intrinsic structure of the peptide and MHC sequences, making the pSA layers less useful than the cSA layers. We still think that our cross-attention architecture can be useful when applied to suitable data.

